# DeepRES: Deep learning enables reaction-based comprehensive enzyme screening

**DOI:** 10.1101/2025.07.28.667344

**Authors:** Keisuke Hirota, Takuji Yamada

## Abstract

**Background:** Enzymes accelerate biochemical reactions in living organisms, thus playing an important role in metabolism. Although metabolic pathway databases are growing, many metabolic reactions, termed orphan enzymes, have not been annotated to gene sequences, which hinders functional annotation in genomic analysis. Moreover, protein databases contain many proteins of unknown function. Owing to this gap between known proteins and enzymatic reactions, various proteins of unknown function may be orphan enzymes; however, available tools cannot adequately predict these links.

**Results:** In this study, we developed DeepRES, an AI-based framework for comprehensive enzyme screening, to explore novel enzyme candidates from proteins of unknown function for reactions of interest. DeepRES implements enzyme screening via two steps: classification of enzymes and non-enzymes and prediction of catalytic capabilities for enzyme‒reaction pairs. The two deep learning models comprising DeepRES showed comparable or superior performance to that of existing software. We performed screening of 1,255 orphan enzymes involved in the microbiome using DeepRES and successfully identified candidate proteins for 897 orphan enzymes. We then used those candidates as references for genomic analysis and explored novel biosynthetic gene clusters from microbial genomes to obtain promising candidate gene clusters, including those related to anthocyanin degradation.

**Conclusions:** Comprehensive enzyme screening via DeepRES, which is the first computational tool designed to associate orphan enzymes with proteins of unknown function, is expected to facilitate high-throughput identification of orphan enzyme-encoding genes. Furthermore, DeepRES can be easily integrated into the current genomic analysis pipeline to extend the functional annotation.

## Background

Enzymes are proteins that exhibit catalytic activity and accelerate biochemical reactions in living organisms. A series of enzymatic reactions is referred to as a metabolic pathway, and metabolic pathway databases, such as the Kyoto Encyclopedia of Genes and Genomes (KEGG) [1], are frequently used to infer metabolic function from genome sequences. Despite the accumulation of vast amounts of gene sequence data due to advances in sequencing technologies and analytical software, from approximately 20% to 50% of the reactions in pathway databases remain unassigned to a corresponding gene sequence [1–3]. Enzymes for which gene sequence information related to confirmed activities is lacking are defined as orphan enzymes [4,5]; this lack of sequence information hinders functional analysis, such as functional annotation and metabolic modeling. Therefore, the identification of genes that encode orphan enzymes is essential for the advancement of genomic analysis.

Moreover, protein databases such as UniProt [6] contain many proteins of unknown function. Owing to the gap between known proteins and enzymatic reactions, various proteins of unknown function are expected to be orphan enzymes. Therefore, enzyme screening for proteins of unknown function is necessary for the identification of orphan enzymes. As experimentally verifying protein function is time-consuming and labor-intensive, a computational approach is especially needed [7].

Numerous computational tools for enzyme screening have been developed, and they are categorized into two primary types: enzyme classification methods and enzyme prediction methods. Enzyme classification methods take protein sequences as inputs and output functional annotations. These include homology search tools, such as BLAST [8], and machine learning (ML) models that predict functional labels for input protein sequences [9–14]. Following the recent success of highly accurate protein 3D structure prediction [15,16], structure-based computational methods have flourished, with the development of Foldseek [17], which is a fast protein structure search tool, and ML models for predicting enzymatic functions on the basis of protein structures [18,19]. ML models that are independent of reference databases are helpful in the exploration of orphan enzyme-encoding genes. However, existing models are designed to predict functional classifications such as EC numbers and thus have the limitation that they can handle only enzymatic reactions within the range covered by the classification schema. In our previous work, we developed an ML-based framework that covers a broader range of enzymatic reactions by employing reaction classifications on the basis of the chemical structural transformation patterns of substrates and products; however, it still cannot handle all reactions owing to its dependence on the reaction classification schema [3]. Furthermore, most existing approaches have limitations in the prediction of enzyme classification for non-enzymes as they do not consider the input of non-enzymes. In contrast, enzyme prediction methods such as E-zyme 2.0 [20] take any reaction as input and output enzymes that catalyze similar reactions. While these approaches can handle all reactions, including orphan enzyme reactions, their search space is restricted to known enzymes; they cannot search for orphan enzyme candidates from proteins of unknown function. Most existing computational tools do not adequately predict whether a protein of unknown function is likely to be an orphan enzyme.

Implementing a computational framework that enables direct and transverse searches for proteins and enzyme reactions allows for the exploration of enzyme candidates from all proteins, including those with unknown functions, for any given reaction. To this end, multi-modal AI that processes proteins and reactions has the potential to achieve this goal, and cross-modal retrieval via multi-modal AI is expected to be a compelling avenue. Originally introduced within the computer-vision community, contrastive language–image pre-training (CLIP) [21] aligns natural-language captions with their corresponding images by projecting both modalities into a shared latent space; embedding proteins and enzymatic reactions into a shared latent space, as in CLIP, would enable comprehensive enzyme screening as a cross-modal retrieval.

In this study, we developed DeepRES, which is an AI-based framework for comprehensive enzyme screening, to explore novel enzyme candidates from proteins of unknown function for reactions of interest. DeepRES consists of a protein classification model (EnzymeCNN) and an enzyme screening model (EnzymeCLIP) (Figure 1A). We show that EnzymeCNN and EnzymeCLIP achieve comprehensive enzyme screening and outperform existing computational tools. We subsequently focused on human gut bacteria because of the abundance of orphan enzymes and genes of unknown function [2,22], and the availability of extensive metabolic pathway and protein 3D structure resources [2,23], therefore we explored candidate proteins from the ESM Metagenomic Atlas [23] for 1,255 orphan enzymes.

**Figure 1:**
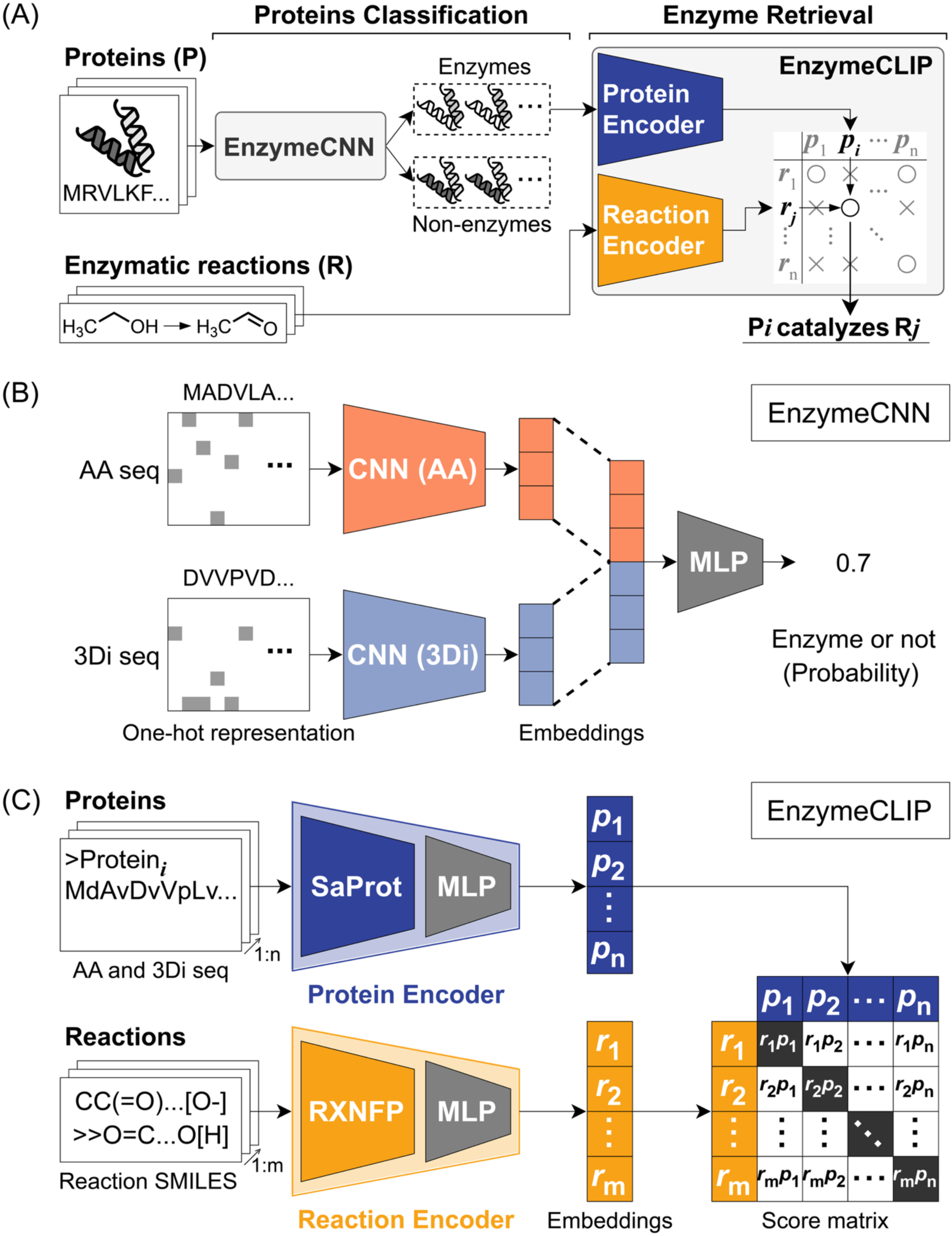
Overview of DeepRES. (A) DeepRES consists of two deep learning models: EnzymeCNN and EnzymeCLIP. First, DeepRES takes proteins as inputs and classifies them into enzymes and non-enzymes via EnzymeCNN. Next, DeepRES associates the predicted enzymes with input reactions by EnzymeCLIP. (B) EnzymeCNN is a convolutional neural network (CNN)-based model for protein classification. EnzymeCNN classifies input proteins into enzymes and non-enzymes by processing amino acid sequences and 3Di sequences separately using CNNs and integrating the sequence and structural information. (C) EnzymeCLIP is a contrastive language–image pre-training (CLIP)-like model for enzyme retrieval. EnzymeCLIP is based on two pretrained language models, SaProt and RXNFP, and embeds input proteins and reactions into a shared latent space by the protein encoder and the reaction encoder. Thus, EnzymeCLIP can comprehensively predict the enzymatic activities of enzyme-reaction pairs of interest. Proteins in structure-aware sequences and enzymatic reactions in Reaction SMILES notation are passed to EnzymeCLIP.

## Results

### DeepRES framework specification

DeepRES is an AI-based framework for comprehensive enzyme screening that comprises two deep learning models: EnzymeCNN and EnzymeCLIP (Figure 1A). EnzymeCNN is a convolutional neural network (CNN)-based model used to classify enzymes and non-enzymes according to protein sequence and structure (Figure 1B). EnzymeCLIP is a CLIP-like model used to predict enzymatic activities for protein‒reaction pairs by projecting proteins and reactions into the same embedding space and calculating cosine similarities as scores (Figure 1C). DeepRES performs enzyme screening by taking proteins and reactions as inputs. DeepRES first classifies proteins as enzymes or non-enzymes via EnzymeCNN. For those proteins predicted to be enzymes, EnzymeCLIP then predicts their correspondence to given reactions. These processes enable DeepRES to consider that input proteins might be non-enzymes and to be independent of the reaction classification schema.

### EnzymeCNN development

#### EnzymeCNN effectively integrates protein sequence and structural information

We performed hyperparameter optimization of EnzymeCNN on a development dataset constructed from the Swiss-Prot [6] and the AlphaFold Protein Structure Database (AlphaFold DB) [15]. The results demonstrated that EnzymeCNN is robust to its hyperparameters and can classify enzymes and non-enzymes accurately (Table 1, Supplementary Figure S3). To isolate the contribution of each protein modality, we constructed two ablation models: EnzymeCNN-AA, which receives only amino acid sequences, and EnzymeCNN-3Di, which receives only 3Di sequences that represent protein 3D structures. After both ablation models were optimized via the same procedure as that for EnzymeCNN, the best EnzymeCNN model outperformed both ablations, which suggests the effectiveness of integrating protein sequences and structural information in the protein classification task (Figure 2, Supplementary Figure S3). A comparison of EnzymeCNN-AA and EnzymeCNN-3Di indicated that protein 3D structures mainly help in classifying enzymes and non-enzymes.

**Table 1:** Results of EnzymeCNN hyperparameter tuning.

| Number of ResNet layers | Embedding dim | Accuracy | F1-score | MCC | AUC |
| --- | --- | --- | --- | --- | --- |
| 3 | 256 | 0.9671 | 0.9664 | 0.9342 | 0.9920 |
| 3 | 512 | 0.9671 | 0.9664 | 0.9343 | 0.9918 |
| 4 | 256 | 0.9696 | 0.9690 | 0.9393 | 0.9926 |
| 4 | 512 | 0.9689 | 0.9683 | 0.9378 | 0.9926 |
| 5 | 256 | 0.9722 | 0.9717 | 0.9444 | 0.9933 |
| 5 | 512 | 0.9692 | 0.9685 | 0.9383 | 0.9927 |

**Figure 2:**
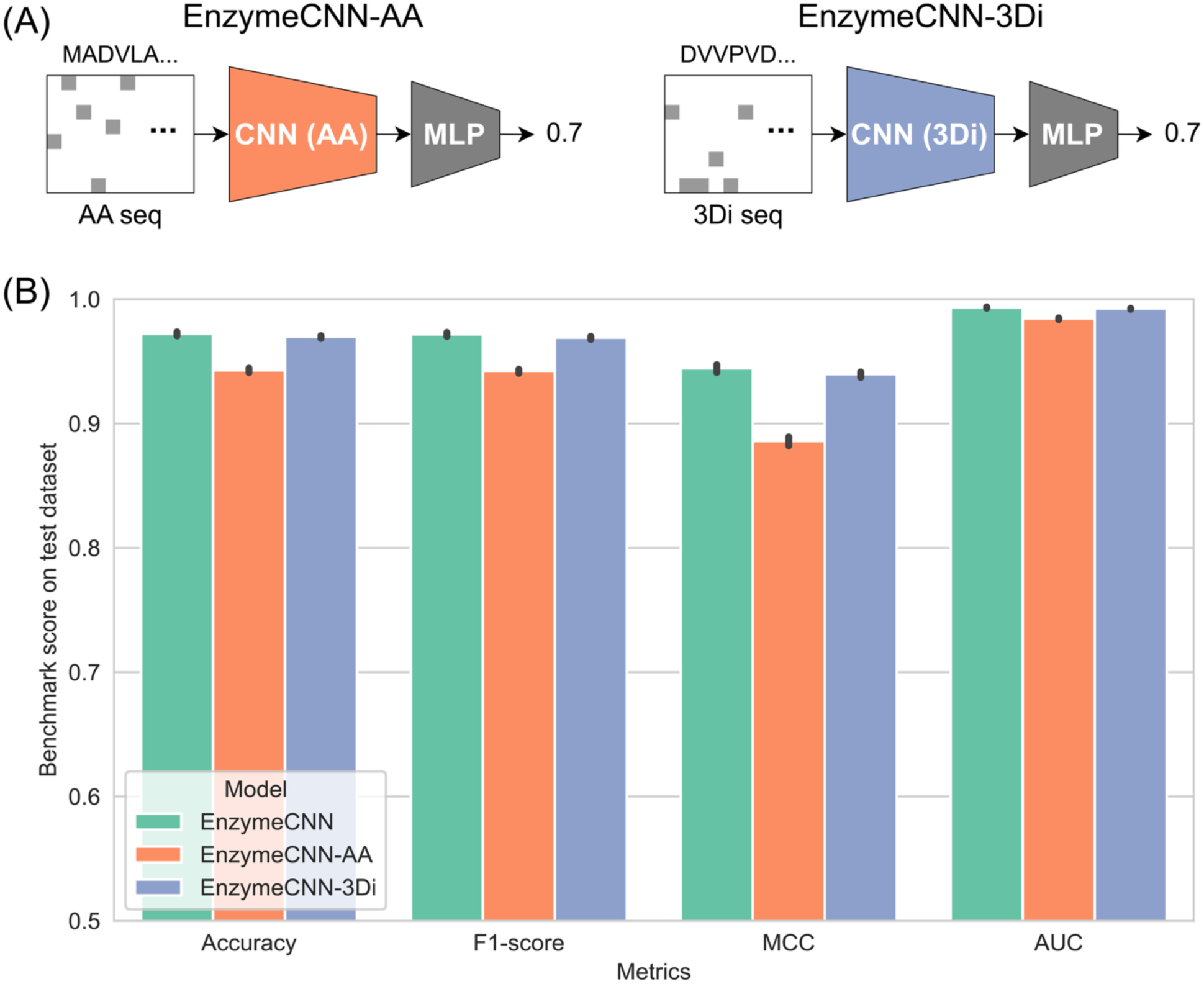
Performance comparison of EnzymeCNN with ablation models. (A) The two ablation models are EnzymeCNN-AA, which receives only amino acid sequences, and EnzymeCNN-3Di, which receives only 3Di sequences. (B) EnzymeCNN was compared with EnzymeCNN-AA and EnzymeCNN-3Di in terms of performance on the test dataset after hyperparameter tuning.

#### EnzymeCNN outperforms existing algorithms in the protein classification task

ML models are generally more sensitive to the functional similarity of proteins than are homology search algorithms [9,10]. Thus, we evaluated the performance of EnzymeCNN on proteins whose sequences or structures are far from the training data to assess whether EnzymeCNN is better at detecting functional similarity than existing search methods are. Specifically, we removed validation and test data similar to training data in protein sequences or structures using MMseqs2 [24] and Foldseek [17], re-optimized the EnzymeCNN hyperparameters and compared EnzymeCNN with MMseqs2 and Foldseek in the protein classification task (Supplementary Tables S1 and S2). Across all similarity thresholds, EnzymeCNN outperformed the existing tools (Figure 3). Notably, when the similarity threshold was tightened, the performance of MMseqs2 and Foldseek collapsed, whereas EnzymeCNN retained its performance.

**Figure 3:**
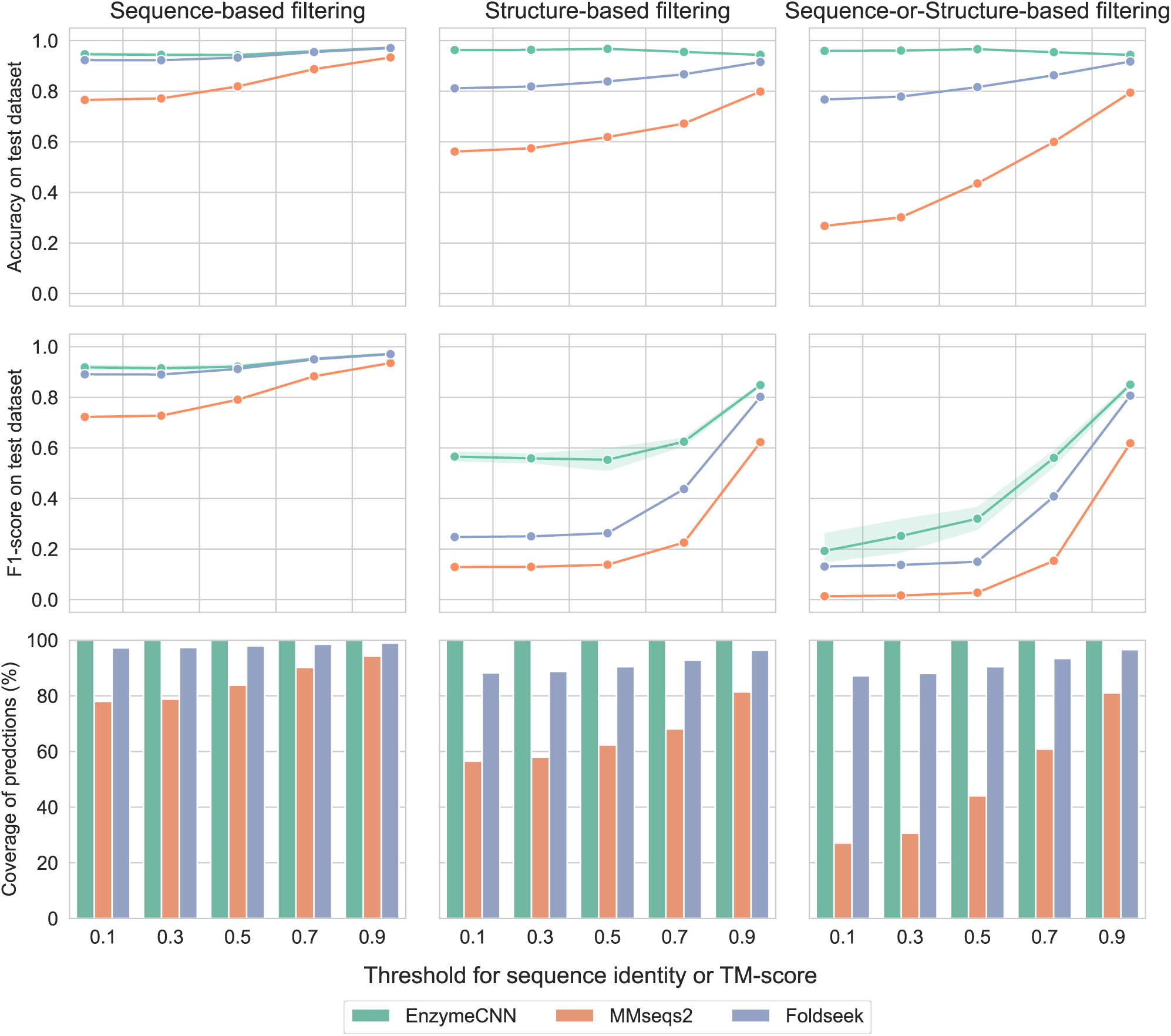
Performance comparison of EnzymeCNN with existing tools under protein similarity-limited situations. After removing validation and test data similar to the training data in terms of sequence identity and/or TM-score via MMseqs2 and/or Foldseek, EnzymeCNN and the two previous approaches were evaluated on the protein similarity-limited dataset. The bottom row shows the percentage of the test data that each tool was able to annotate as enzymes or non-enzymes. When MMseqs2 and Foldseek were evaluated, no hits were treated as incorrect.

The advantage of EnzymeCNN persisted when all computational approaches were benchmarked on new Swiss-Prot data. Compared with both MMseqs2 and Foldseek, EnzymeCNN consistently achieved higher overall accuracy and F1-scores (Figure 4). In addition, unlike the homology search, EnzymeCNN annotated all input proteins as enzymes or non-enzymes.

**Figure 4:**
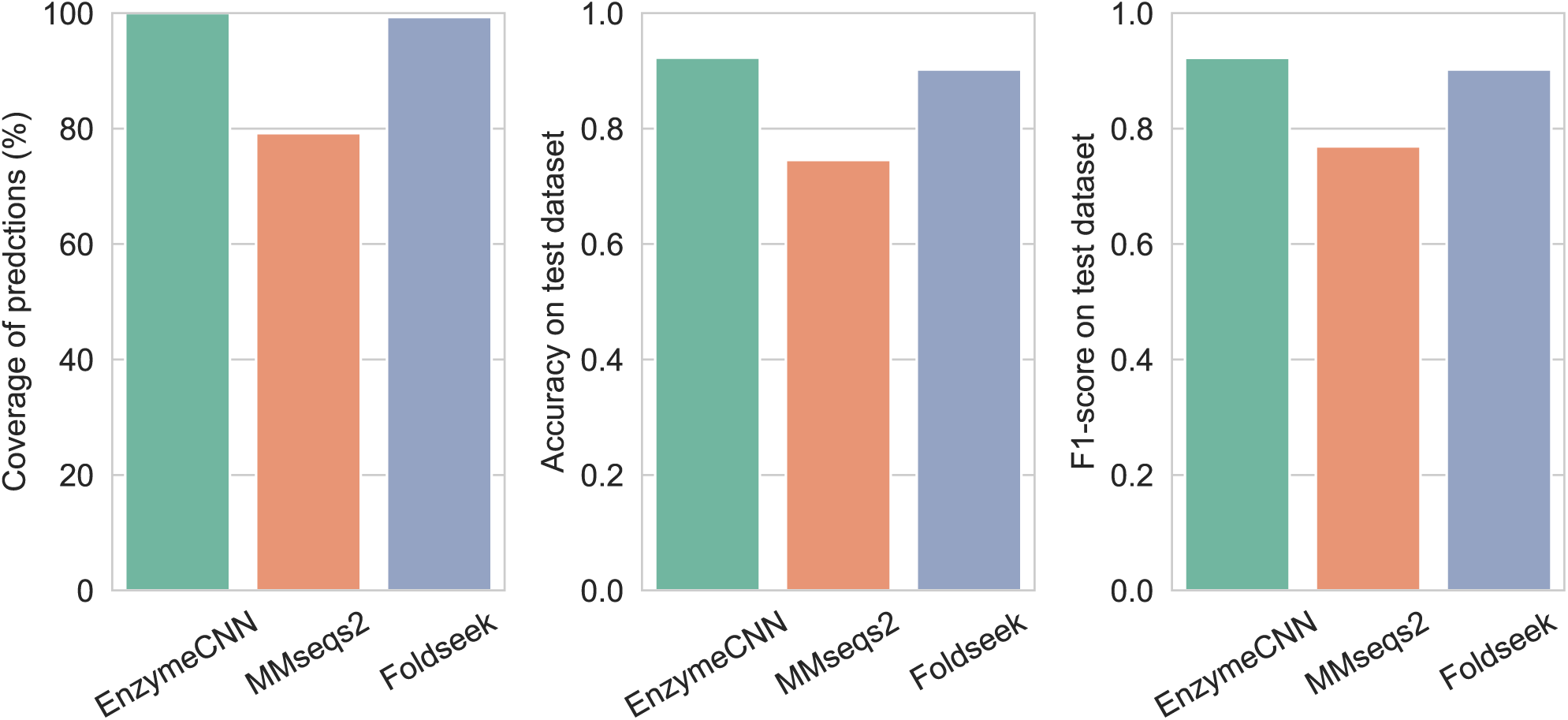
Performance comparison of EnzymeCNN with existing tools on the new protein dataset. EnzymeCNN, MMseqs2 and Foldseek were evaluated on the new protein dataset. The left graph shows the percentage of new data that each tool was able to annotate as enzymes or non-enzymes. In evaluating MMseqs2 and Foldseek, no-hits were treated as incorrect.

### EnzymeCLIP development

#### EnzymeCLIP performed well in the enzyme retrieval task

Using a development dataset assembled from Swiss-Prot [6], AlphaFold DB [15] and Rhea [25], we systematically tuned the architecture of EnzymeCLIP and its training settings, such as loss functions, batch sizes and fine-tuning of the two pretrained models (Supplementary Table S3). The experimental results demonstrated that fine-tuning the underlying language models in EnzymeCLIP is effective for enhancing performance. SaProt-based models consistently outperformed ESM-2-based models, thus underscoring the importance of integrating protein sequence and structural information in the enzyme retrieval task. In contrast, the performance of EnzymeCLIP was largely insensitive to the choice of batch size across the range we tested. The use of cyclic loss improved the performance of EnzymeCLIP, whereas replacing contrastive loss with soft contrastive loss led to a decline in performance.

#### EnzymeCLIP outperformed the state-of-the-art method in the EC number prediction task

Since another computational approach similar to EnzymeCLIP has not yet been established, we compared EnzymeCLIP with CLEAN [9], which is the state-of-the-art enzyme function annotation tool, in the EC number prediction task on new Swiss-Prot and Rhea data (Supplementary Figure S4). Although CLEAN is a multi-label classifier and EnzymeCLIP is designed for a screening task, EnzymeCLIP performed competitively with CLEAN in terms of top-1 accuracy and outperformed CLEAN in terms of top-10 accuracy (Figure 5).

**Figure 5:**
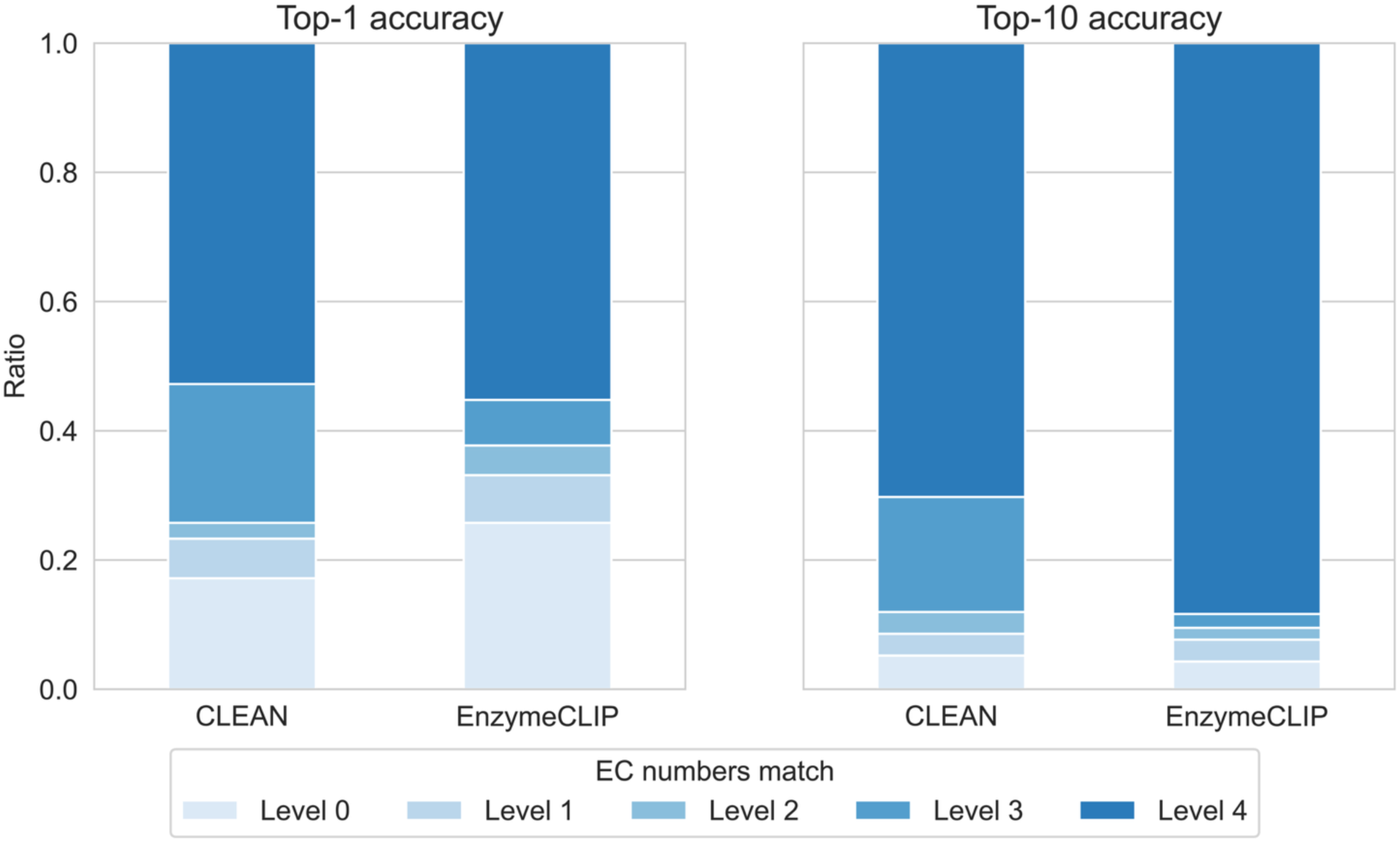
Performance comparison of EnzymeCNN with CLEAN in the EC number prediction task. Performance comparison of EnzymeCNN with CLEAN on the new enzyme‒reaction dataset in terms of the top-1 accuracy and the top-10 accuracy. If the actual EC number is 1.1.1.1 and the predicted EC number is 1.1.2.3, the EC numbers match at level 2.

### Application of DeepRES to metagenome data to explore orphan enzyme candidates

We focused on metagenome data and proteins of unknown function associated with orphan enzymes using DeepRES. Specifically, we applied DeepRES to 13,902,058 metagenomic proteins of unknown function obtained from the ESM Metagenomic Atlas [23] and 1,255 orphan enzymes involved in human gut bacteria recorded in EnteroPathway [2] (Figure 6). First, the 13,902,058 proteins were classified as enzymes and non-enzymes by EnzymeCNN, which resulted in 595,340 predicted enzymes. Next, we explored candidate proteins for the 1,255 orphan enzymes identified by EnzymeCLIP, and associated 97,074 proteins of unknown function with 897 orphan enzymes. Furthermore, to annotate the remaining predicted enzymes with enzymatic reactions, we explored candidate proteins for well-known enzymatic reactions in Rhea (release 138) using EnzymeCLIP (Supplementary Figure S5).

**Figure 6:**
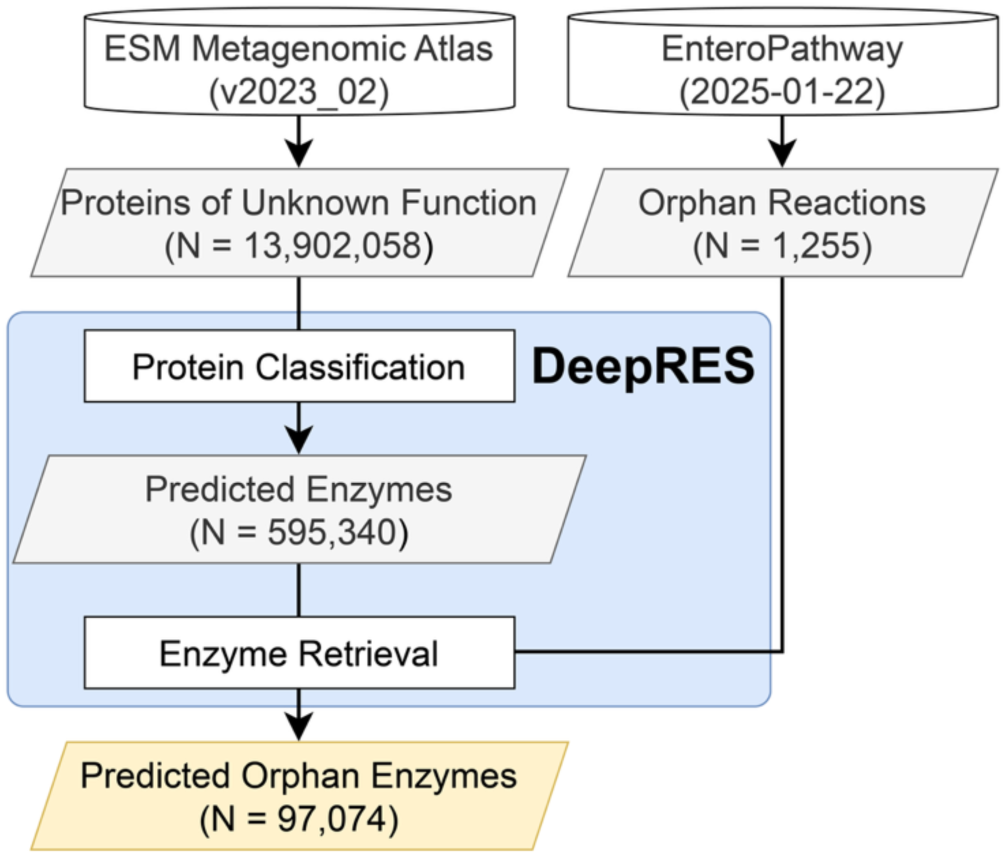
Flow and results of the application of DeepRES to metagenome data.

We obtained 4,744 representative metagenome-assembled genomes (MAGs) of human gut bacteria from MGnify (Unified Human Gastrointestinal Genome v2.0.1)[26] and queried all genes in the MAGs against the orphan enzyme candidates identified by DeepRES as reference databases using BLAST. The homology search against putative orphan enzyme proteins enabled us to assign the functional annotations of the orphan enzymes to the MAGs. Inspired by previous studies [3,27], we attempted to identify novel biosynthetic gene clusters (BGCs) that correspond to metabolically related orphan enzymes by evaluating orphan enzyme candidates in microbial genomes, with consideration of genomic and metabolomic neighboring information (Table 2). For example, BGC screening for EPM0520 and EPM0522 related to anthocyanin degradation (Figure 7) identified novel BGC candidates from 50 and 98 microbial genomes, respectively (Supplementary Table S4).

**Figure 7:**
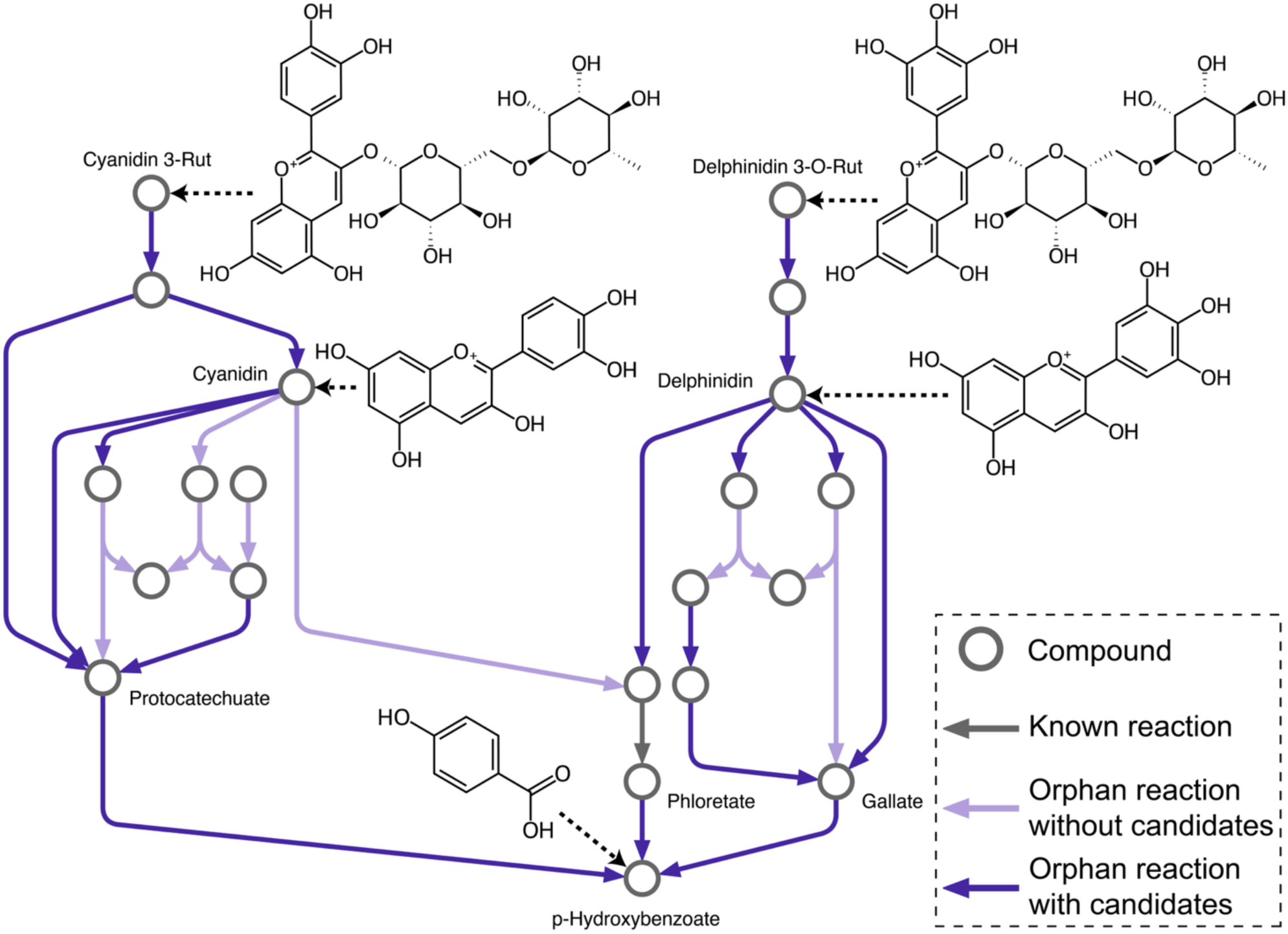
Examples of BGC screening with putative orphan enzyme proteins obtained by DeepRES. The two anthocyanin degradation modules with the most orphan enzymes are EPM0520 (cyanidin-3-rutinoside degradation) and EPM0522 (delphinidin 3-O-rutinoside degradation).

**Table 2:**
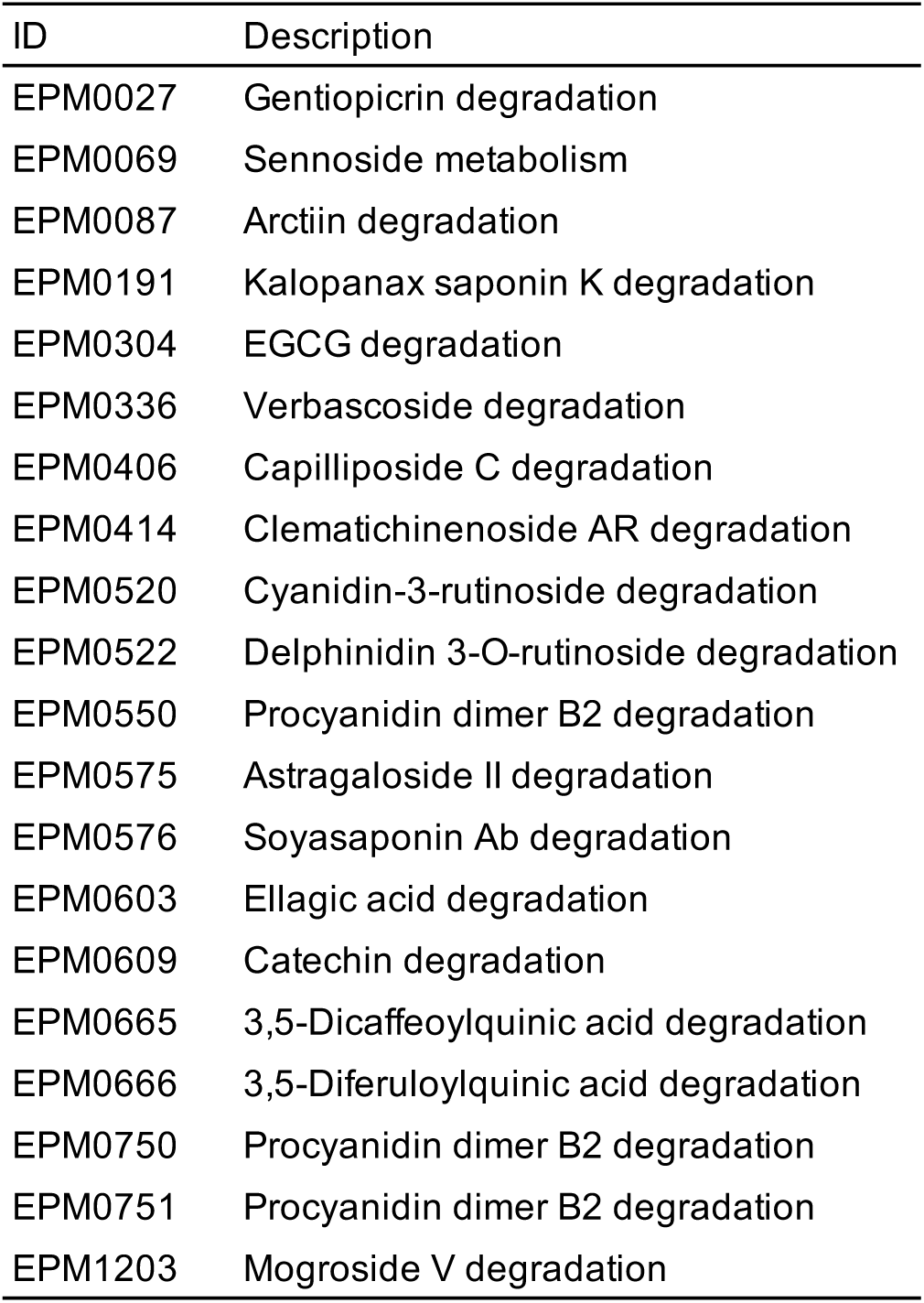
List of modules subjected to a novel biosynthetic gene cluster search.

## Discussions

Metabolic pathway databases are continually growing due to the accumulation of results from previous studies, and many enzymatic reactions in the databases remain unassigned to corresponding gene sequences; the associated enzymes are referred to as orphan enzymes. In this study, which focused on the possibility that some proteins of unknown function are orphan enzymes, we developed DeepRES, which is an AI-based framework for comprehensive enzyme screening, to explore novel enzyme candidates from proteins of unknown function for reactions of interest. To our knowledge, DeepRES is the first method designed to associate any reaction, including those that involve orphan enzymes, with any protein, including functionally unknown proteins. The two models that comprise DeepRES outperformed existing computational tools. We applied DeepRES to metagenome data, which consisted of 13,902,058 metagenomic proteins of unknown function and 1,255 orphan enzymes involved in the metabolism of human gut bacteria. We classified the 13,902,058 proteins into enzymes and non-enzymes via EnzymeCNN and successfully mapped the 97,074 predicted enzymes to 897 orphan enzymes using EnzymeCLIP.

In addition to its primary role in comprehensive enzyme screening, DeepRES provides a catalog of putative enzymes for homology search. The use of the orphan enzyme candidates identified by DeepRES as a reference database is also beneficial because this approach does not require a GPU and can be easily integrated into the current genomic analysis pipeline, particularly for functional annotation. We demonstrated that this approach enables us to identify orphan enzyme candidate genes that are linked to a taxonomy and a genomic location within microbial genomes. Furthermore, following the findings of previous studies that integrating genomic and metabolomic neighboring information facilitates the identification of orphan enzyme-encoding genes [3,27], we successfully identified BGC candidates from microbial genomes for 15 EnteroPathway modules, which were composed mostly of orphan enzymes (Supplementary Table S4). These modules included anthocyanin degradation (EPM0520 and EPM0522). Anthocyanins, such as cyanidin and delphinidin have been suggested to act on obesity, type 2 diabetes, inflammatory bowel disease and colorectal cancer through their conversion into various metabolites by the human gut microbiota, thus they have attracted attention for dietary therapy [28,29]. Despite its high importance, the mechanisms of anthocyanin degradation in the human gut microbiota remain largely unclear [1,2]. Novel BGC candidates were detected for EPM0520 and EPM0522 in 50 and 98 microbial genomes, respectively, primarily in *Bacteroides* (Supplementary Table S4). These results are promising because *Bacteroides* is capable of metabolizing anthocyanin [28,29]. In addition, more than 80% of the bacteria with BGC candidates involved in anthocyanin degradation belong to *Bacteroidota*, which suggests a potential association between the anthocyanin-induced decrease in the *Firmicutes*/*Bacteroidetes* ratio in the gut microbiota [28,30] and the novel BGC candidates identified in this study. Moreover, although DeepRES was used for microbiome research in this study, DeepRES is applicable to eukaryotes and viruses.

While DeepRES successfully mapped many previously uncharacterized proteins to specific enzymatic reactions, including orphan enzymes, approximately 40% of the proteins predicted by EnzymeCNN to be enzymes were not linked to enzymatic reactions (Supplementary Figure S5). This result arose from the nature of EnzymeCLIP which uses a reaction dataset as a reference database, and indicates the limitations of DeepRES’s applicability, as well as the possibility that the remaining predicted enzymes might catalyze unknown enzymatic reactions.

Unlike existing approaches, DeepRES is independent of reaction classification, and its prediction can be converted into any preferred classification schema, as demonstrated in its comparison with CLEAN in the EC number prediction task. However, there is room for improvement in the numerical representation of enzymatic reactions in DeepRES. DeepRES is based on RXNFP, which embeds Reaction SMILES representation via natural-language processing (NLP) techniques; however, the NLP models pose difficulties in recognizing chirality [31]. Given that enzymatic reactions are characterized by high stereoselectivity, improving the underlying reaction encoding is expected to further increase the effectiveness of DeepRES.

Although increasing the batch size is generally effective in contrastive learning [21,32,33], this trend was not observed during the development of EnzymeCLIP. The multimodal data of proteins and enzymatic reactions include many-to-many relationships due to homologous proteins, protein multifunctionality and convergent evolution. These complex associations may offset the benefit of larger batches by introducing many-to-many relationships within mini-batches that CLIP cannot account for. That is, the results suggest the limitations of a CLIP-like approach for biological multi-modal data. Inspired by SoftCLIP [34], we introduced a soft contrastive loss to relax the assumptions regarding one-to-one relationships in the CLIP strategy; however, this approach decreased the performance of EnzymeCLIP. This could be explained by the possibility that the global similarity of proteins obscures the similarities of their enzymatic functions. Taken together, CLIP is applicable to any multi-modal data, thus enabling comprehensive enzyme screening in this study; however, its application to biological multi-modal data still leaves room for improvement.

## Conclusions

We developed DeepRES, which is a deep learning-based computational tool for comprehensive enzyme screening. Key features that distinguish DeepRES from previous approaches are the explicit consideration of proteins of unknown function and its independence of reaction classification schema, such as EC numbers. DeepRES identified candidate proteins for 897 orphan enzymes associated with human gut bacteria. Furthermore, we explored novel gene clusters involved in human-gut interactions by employing those candidates as reference datasets in a homology search. The results demonstrate the potential of DeepRES to bridge the gap between our understanding of gene sequences and functions and to expand the current genome analysis. DeepRES is expected to contribute to the high-throughput and cost-effective identification of genes that encode orphan enzymes, thereby promoting a deeper understanding of metabolism and advancing enzyme engineering.

## Methods

### Dataset creation

To develop DeepRES, we constructed a protein dataset for protein classification and an enzyme‒reaction dataset for enzyme screening. All the data were retrieved from UniProt [6], AlphaFold DB [15] and Rhea [25]. First, all Swiss-Prot protein sequences were obtained from UniProt (release 2024_03) and subjected to homology clustering using MMseqs2 (v15-6f452) [24] with an identity threshold of 0.9, which yielded 335,684 cluster representatives. Predicted protein 3D structure data from Swiss-Prot were retrieved from AlphaFold DB (v4), and those data were converted to 3Di sequences using Foldseek (v9-427df8a) [17]. The 21,520 cluster representatives that lacked 3Di sequences were discarded. The 19,642 cluster representatives longer than 850 residues were also discarded because of computational constraints. Amino acid and 3Di sequences were then merged for each protein, which resulted in 294,522 protein sequences.

Next, the protein dataset was split 8:1:1 into training, validation, and test datasets, with a consistent ratio of enzymes to non-enzymes ensured for each split. This resulted in 235,617 training data points, 29,452 validation data points, and 29,453 test data points. The of enzyme and non-enzyme labels were assigned according to the presence and absence, respectively, of EC number annotations in Swiss-Prot; enzymes accounted for approximately 49% of each split.

Finally, the protein dataset was divided into enzymes and non-enzymes, and the enzymes were mapped to enzymatic reactions obtained from Rhea (release 133) to construct an enzyme‒reaction dataset that contained 139,156 enzyme‒reaction pairs. Since proteins and reactions have many-to-many relationships due to protein multifunctionality and convergent evolution, the enzyme‒reaction dataset was first split at a ratio of 8:1:1 into training, validation and test datasets based on the associated reactions. Then, various training and validation data were removed to avoid overlap in proteins between data splits. This resulted in 106,174 training data points, 14,113 validation data points, and 18,869 test data points.

In addition, to compare the performance of DeepRES with those of previous computational tools, we constructed a new dataset that was not used in their development. Specifically, we utilized Swiss-Prot data updated from version 2024_03 to 2024_06 and Rhea data updated from release 133 to 136 to prepare new protein and enzyme‒reaction datasets. As the 3D structures of most of the new proteins are not yet included in the AlphaFold DB, we performed protein 3D structure prediction via AlphaFold2 [15].

### EnzymeCNN development

EnzymeCNN is a CNN-based model for classifying enzymes and non-enzymes based on protein sequences and structures, which are represented as amino acid and 3Di sequences. As CNNs have proven versatile for sequential data, including text, speech and protein sequences [10,35], we applied CNNs for protein classification especially by utilizing dilated convolution techniques [36], to broaden the receptive field exponentially without losing resolution. Hyperparameter tuning and performance tests were conducted on the development protein dataset five times with different seeds. Each model was trained for 50 epochs, and the checkpoint that yielded the highest validation accuracy was subsequently evaluated on the hold-out test dataset. To validate the benefits of EnzymeCNN approach, which integrates sequence and structural information, we built two ablation variants, EnzymeCNN-AA (sequence only) and EnzymeCNN-3Di (structure only), and tested them like we tested EnzymeCNN.

We next tested the performance of EnzymeCNN under limited sequence or structural homology. Specifically, we removed training data that were similar to validation or test data via homology searches with MMseqs2 and Foldseek, then re-optimized the hyperparameters and evaluated the performance of EnzymeCNN. The searches were performed with the sequence identity or TM-score thresholds set to 0.1, 0.3, …, 0.9 and the other parameters set to the MMseqs2 default values. For benchmarking, we additionally classified enzymes and non-enzymes by querying the test data against the training dataset using MMseqs2 and Foldseek with default settings. Finally, we evaluated the performance of EnzymeCNN on the new protein dataset. The development dataset was split 9:1 into training and validation datasets, which were used to build EnzymeCNN, MMseqs2 and Foldseek were again used as benchmark methods.

### EnzymeCLIP development

EnzymeCLIP is a CLIP-like model for comprehensive enzyme screening built on two pretrained, modality-specific language models̶SaProt [37], which is a structure-aware protein language model, and RXNFP [38], which is a BERT-based encoder for chemical reactions. EnzymeCLIP takes protein sequence and structural information, as well as reaction information, as inputs to predict catalytic activity for all combinations of input proteins and reactions. Protein information is supplied to EnzymeCLIP in the “structure-aware vocabulary” format introduced with SaProt, which integrates primary- and tertiary-structure features into a unified text representation. Reaction information is provided in Reaction SMILES format, which is a textual representation of chemical reactions. EnzymeCLIP embeds proteins and reactions into a latent space using the two separate encoders and is trained to pull vector representations of an enzymatic reaction and its catalytic protein close and to push those of non-catalytic pairs apart. Consequently, catalytic activity can be inferred by computing the cosine similarity between the resulting vectors. Thus, the catalytic activities of protein‒reaction pairs can be predicted by calculating cosine similarities between the embedding vectors of proteins and reactions.

To improve the performance of EnzymeCLIP, both SaProt and RXNFP are fine-tuned within EnzymeCLIP using Low-Rank Adaptation (LoRA) [39]; the hyperparameters of LoRA are the same as those used in previous studies [39,40]. To assess the contribution of each element in EnzymeCLIP, we constructed ablation variants that (i) replace SaProt with ESM-2 [23], (ii) omit the fine-tuning of pretrained language models and (iii) vary the batch size during training. We further examined loss functions inspired by previous studies [34,41]: contrastive loss, soft contrastive loss and cyclic loss. These loss functions are computed on the basis of a batch of *N* protein‒reaction pairs 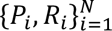 where *P_i_* represents a protein and *R_i_* represents a reaction. EnzymeCLIP converts *P_i_* and *R_i_* into a protein embedding 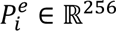 and a reaction embedding 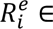 ℝ^256^, respectively. A protein-reaction similarity vector ***p****_i_*(*P*, *R*) and a reaction-protein similarity vector ***p****_i_*(*R*, *P*) are calculated as follows:

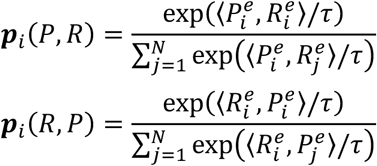

where 〈·,·〉 represents the inner product and where τ represents a trainable temperature parameter. The CLIP strategy uses the one-hot vector that corresponds to the input pairs 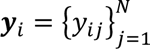 as labels; therefore, the contrastive loss ℒ_CLIP_ is computed as follows:

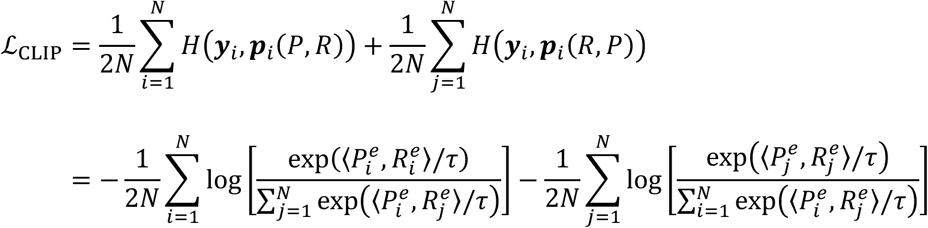

We utilized the similarity between proteins ***p****_i_*(*P*, *P*) to soften the one-hot label vector 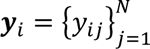 and calculate the soft contrastive loss ℒ_soft_ as follows:

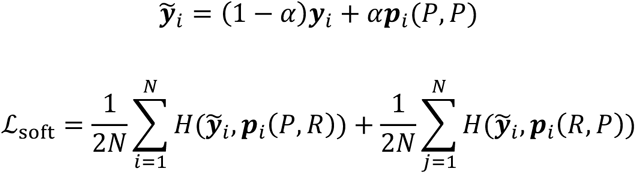

where α is a hyperparameter, which is set to 0.3. Moreover, to align the shared embedding space for each modality in EnzymeCLIP, we introduce the cyclic loss ℒ_cyclic_, via following regularization term:

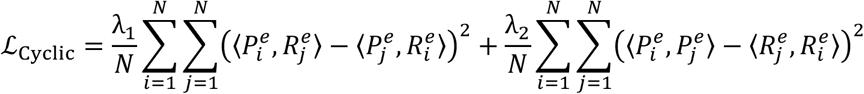

where λ_1_ and λ_2_ are the hyperparameters, which are set to 0.25.

In total, 48 enzyme screening models (the original EnzymeCLIP and 47 variants) were trained for 10 epochs on the development enzyme‒reaction dataset 5 times with different seeds. Their performance was evaluated in terms of the enrichment factor (EF) [42], which is a metric commonly used in screening tasks.

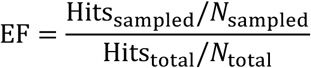

EF measures how concentrated positives are within highly ranked predictions. For example, EF_0.10_ evaluates the top 10% of predictions and takes values from 1 to 10, with a higher EF indicating better screening. For each model, the checkpoint that achieves the highest validation EF_0.10_ was subsequently evaluated on the hold-out test dataset.

Finally, we evaluated the performance of EnzymeCLIP on the new enzyme‒reaction dataset. To build EnzymeCLIP, the development dataset was split 9:1 into training and validation datasets on the basis of reactions. Then, some of the training data were removed to avoid overlap in proteins between the data splits. Since another computational method similar to EnzymeCLIP has not yet been established, we compared EnzymeCLIP with CLEAN [9], which is the state-of-the-art enzyme function annotation tool, in the EC number prediction task. Specifically, while CLEAN predicts EC numbers directly via multi-label classification, EnzymeCLIP annotates EC numbers indirectly for the input protein by comparing the input protein embedding with those of enzymatic reactions and retrieving the EC numbers associated with the enzymatic reactions mapped to the input protein. Given the conceptual difference between EnzymeCLIP, which is a screening model, and CLEAN, which is a classification model, we evaluated their performance in terms of the top-k accuracy.

### Orphan enzyme exploration

To associate proteins of unknown function with orphan enzymes and explore novel enzyme candidates, we applied DeepRES to metagenome data, including the ESM Metagenomic Atlas [23], which is a resource of metagenomic protein structures predicted by ESMFold, and EnteroPathway [2], which is a curated metabolic pathway database of the human gut microbiota. We retrieved 42,236,858 proteins from the ESM Metagenomic Atlas (v2023_02) via Foldseek and performed a homology search of the metagenomic proteins against UniRef90 (release 2025_01) [6] using Diamond BLASTp (v2.0.9) [43] with default parameters. Afterward, the proteins were treated as functional unknowns when the homology search (i) yielded no hits, (ii) aligned to “uncharacterized” or “hypothetical” proteins, or (iii) aligned to proteins without any description, which resulted in 13,902,058 proteins of unknown function. We also identified 1,255 orphan enzymes that correspond to enzymatic reactions that are not associated with sequence information on the basis of the correspondence of each entry with KEGG or UniProt information provided by EnteroPathway. For each reaction, the molecular structures of all the substrates and products were retrieved and converted to reaction SMILES representations; 1,150 orphan enzymes for which complete reaction SMILES representations could be generated were retained for downstream analysis.

We applied DeepRES to the 13.9M proteins of unknown function and 1,150 orphan enzymes. In addition, for proteins predicted to be enzymes by EnzymeCNN yet not linked to any orphan enzyme, we assigned functional annotations by mapping them to the known enzymatic reactions in Rhea (release 138) [25] using EnzymeCLIP. Here, the top 10% border of scores for positive examples in the test dataset, as determined during EnzymeCLIP development, was employed as the threshold for the EnzymeCLIP score to consider an enzyme‒reaction pair to be linked.

### Exploring novel biosynthetic gene clusters

Novel enzyme candidates identified by DeepRES could be utilized as a reference dataset for functional annotation in genome analysis. Therefore, for each orphan enzyme for which we identified candidate proteins by DeepRES, we mapped all genes in 4,744 representative MAGs of human gut bacteria obtained from MGnify (Unified Human Gastrointestinal Genome v2.0.1) [26] to the reference dataset using Diamond BLASTp (v2.0.9) [43]. Previous studies have demonstrated that leveraging a combination of metabolic pathway adjacency and genomic neighborhood information is effective for identifying orphan enzyme-encoding genes [3,27]. Following this finding, we explored novel BGCs by assessing whether the candidate genes are densely populated in microbial genomes for EnteroPathway modules, which are composed mostly of orphan enzymes. Here, we specifically investigated novel BGC candidates for 20 modules (Table 2) that consisted of more than 10 reactions, more than 90% of which were orphan, and DeepRES identified candidate proteins for more than 50% of the orphan enzymes. The criterion for the novel BGCs was that at least five homologs of the orphan enzymes in the module were included in the range of 10 continuous genes in the genome.

## Declarations

### Ethics approval and consent to participate

Not applicable.

### Consent for publication

Not applicable.

### Availability of data and materials

Source codes for the DeepRES are available on GitHub under the MIT license (https://github.com/yamada-lab/deepres). Model weights (EnzymeCNN and EnzymeCLIP), EnteroPathway reaction information, the homology search results for ESM Metagenomic Atlas against UniRef90, and the results of DeepRES application to ESM Metagenomic Atlas, EnteroPathway and Rhea are accessible through Zenodo (https://doi.org/10.5281/zenodo.16347933).

### Authors’ contributions

K.H. and T.Y. contributed to the conception and design of the study. K.H. carried out the development of the deep learning models, the genomic analyses, and creating the python library. K.H. wrote the manuscript with contributions from T.Y. T.Y. supervised the study. The authors read and approved the final manuscript.

## Acknowledgements

We thank Felix Salim for fruitful discussions. This study was carried out using the TSUBAME4.0 supercomputer at Institute of Science Tokyo.

