## Supplementary_materials for "DeepRES: Deep learning enables reaction-based comprehensive enzyme screening": Suppelementary_materials.pdf

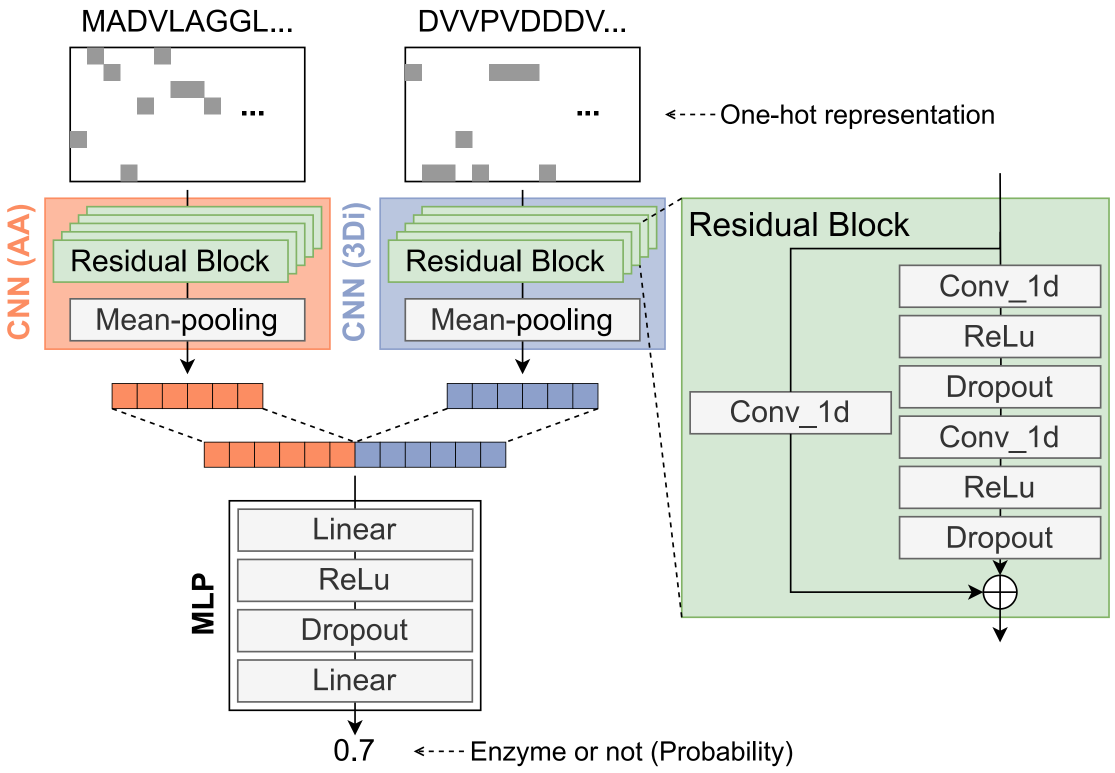


**Figure S1 EnzymeCNN architecture.** EnzymeCNN is a convolutional neural network (CNN)-based model to classify enzymes and non-enzymes according to protein sequence and structure represented as amino acid and 3Di sequences. EnzymeCNN consists of two CNNs and 2-layer multilayer perceptron (MLP). The positional embeddings in the final residual block are collapsed into a single embedding for the input protein by mean-pooling in the CNNs, and then MLP predicts whether the input protein is enzyme or non-enzyme based on the concatenated two embeddings corresponding to an amino acid sequence and a 3Di sequence. In the CNNs, we set filters to 600, dilation rate to 3, kernel size to 9 and batch size to 128. After tuning hyperparameters, the number of residual blocks was chosen to be 5 and the embedding dimension to be 256. To reduce overfitting, we applied weight normalization and dropout at a rate of 0.1 to EnzymeCNN.

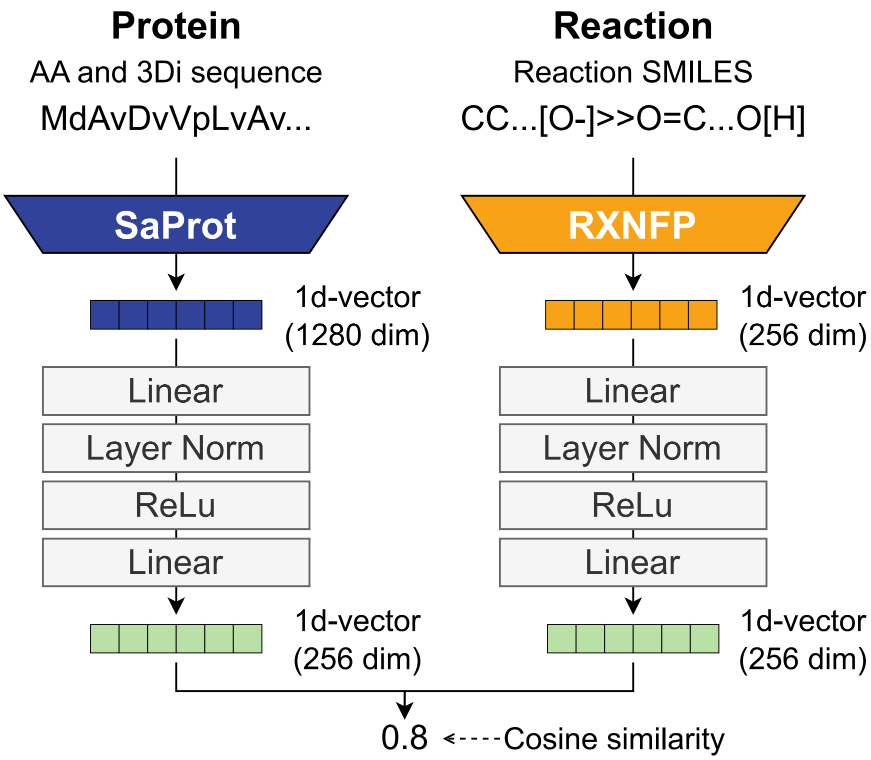


**Figure S2 EnzymeCLIP architecture.** EnzymeCLIP is a contrastive language–image pre-training (CLIP)-like model for comprehensive enzyme screening, builds on two pretrained, modality-specific language models: SaProt and RXNFP. SaProt converts a structure-aware sequence, an integrated representation of an amino acid sequence and a 3Di sequence of a protein, into a 1280-dimensional embedding vector, and RXNFP converts a Reaction SMILES, a text representation of a reaction, into a 256-dimensional embedding vector. EnzymeCLIP learns to project each of those embeddings into a 256-dimensional shared space by 2-layer MLPs, allowing it to predict enzymatic activities for input enzyme-reaction pairs using cosine similarities as scores. We set hidden dimension to 256 in the MLPs and applied layer normalization to the MLPs to reduce overfitting.

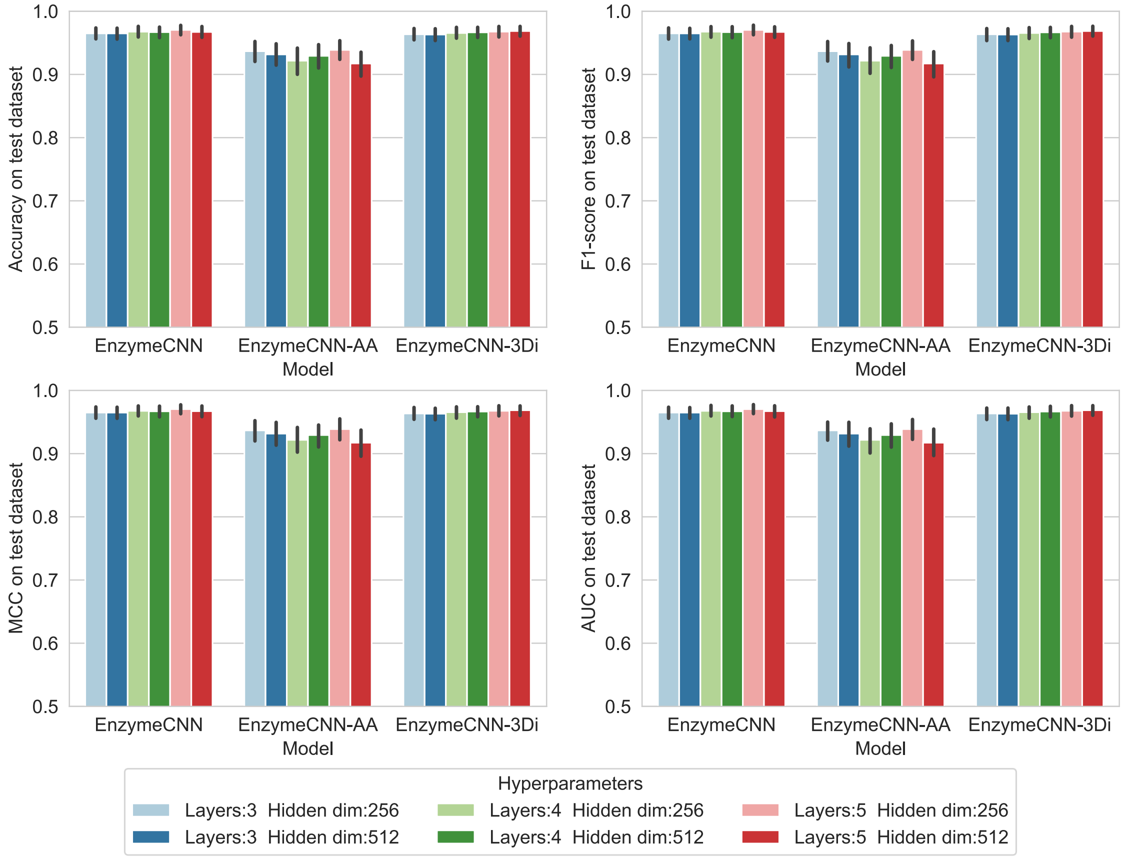


**Figure S3 Hyperparameter tuning of EnzymeCNN and ablation models.**

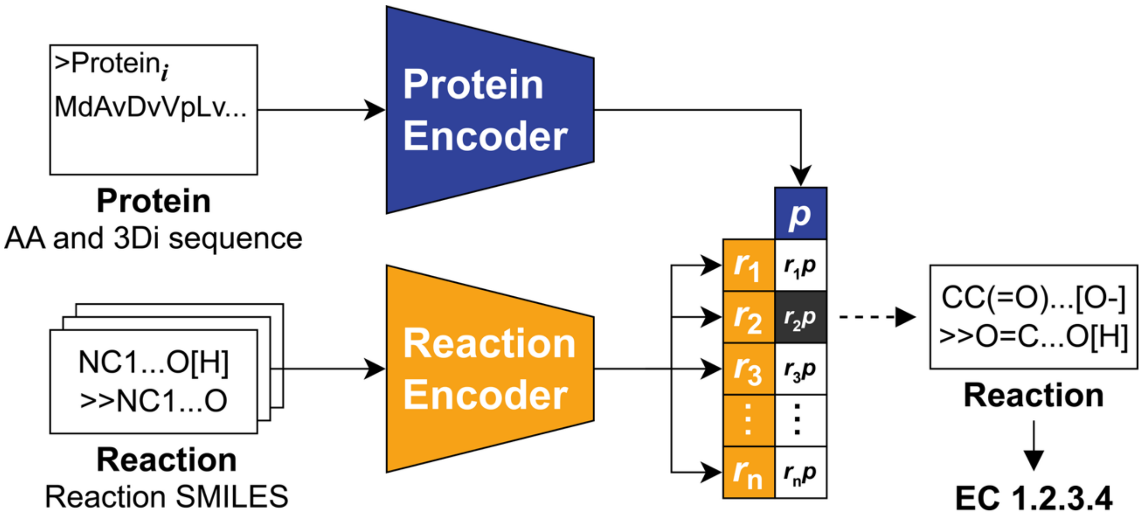


**Figure S4 Flow of EC number prediction using DeepRES.** EnzymeCLIP indirectly annotates EC numbers to the input protein by comparing the input protein embedding with those of enzymatic reactions and retrieving EC numbers associated with the enzymatic reactions mapped to the input protein.

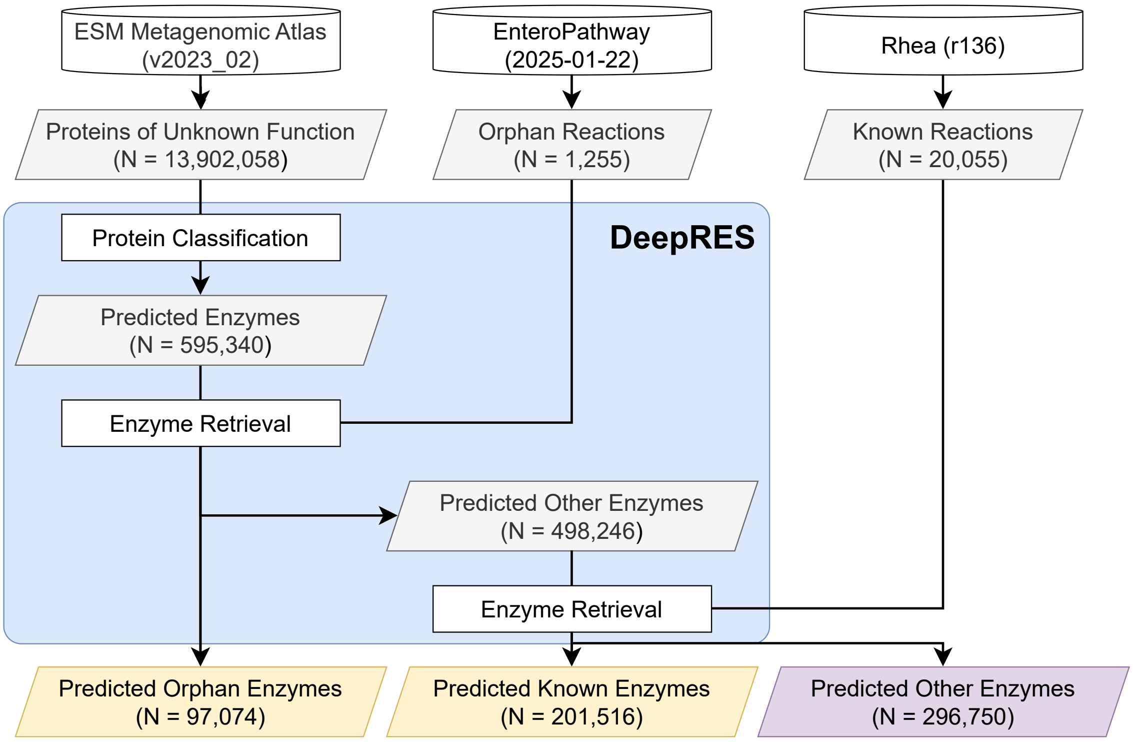


**Figure S5 Reaction-based functional annotation to metagenomic proteins of unknown function using DeepRES.** The flow and result of DeepRES application to metagenomic proteins of unknown function recorded in the ESM Metagenomic Atlas, orphan reactions recorded in EnteroPathway and known reactions recorded in Rhea database.
